# Acute priming using elevated fluid viscosity recovers ‘*young-like*’ single-cell surveillance behaviors in aged human T cells

**DOI:** 10.64898/2026.01.13.699374

**Authors:** Yoseph W. Dance, Alice Amitrano, Annaka M. Saffron, Chanhong Min, Ladaisha T. Thompson, Zhuoxu Ge, Ian M. Smith, Nicolas C. Macaluso, Charles Ezenwanne, Nicholas Milcik, Abigail Fennell, Kendall A. Pyndell, Kimberly M. Stroka, Jeremy D. Walston, Sean X. Sun, Konstantinos Konstantopoulos, Jude M. Phillip

## Abstract

Aging is a complex biological process, often characterized by increased vulnerability to disease, infection, and death. This increased vulnerability is mechanistically linked to a progressive and functional decline of the immune system. In humans, aged lymphocytes lose their capacity to effectively surveil within diverse microenvironments, decreasing their capability for clearing infections and maintaining physiological homeostasis. However, specific mechanisms by which aged lymphocytes, specifically T cells, lose this capacity to surveil remain unclear. We profiled three core characteristics of T cell surveillance at single-cell resolution, specifically migration, deformability, and sensing. While aged T cells retained their capacity for spontaneous migration, they exhibited impaired cellular deformability and deficiencies in sensing local signaling cues. To modulate this surveillance defect, we performed mechanical reprogramming using elevated fluid viscosity. Results showed that acute priming of aged T cells with elevated fluid viscosity recovered a transient *young-like* surveillance phenotype, which was mechanistically linked to membrane tension, cortical F-actin, and Arp3 expression. These findings reveal a key source of surveillance defects in aged T cells and provide an effective mechanical approach to tuning their single-cell behaviors.

**Teaser:** Recovery of ‘young-like’ surveillance phenotypes in aging human T cells via viscosity priming

## INTRODUCTION

Lymphocyte migration and surveillance within complex tissue microenvironments are essential for maintaining physiological homeostasis and regulating overall organismal health^1^. Impaired immune surveillance, particularly in T cells, is closely linked to several aging-related dysfunctions and disease states including cancers, chronic infections, type II diabetes, and rheumatoid arthritis^2–6^. Although impaired surveillance is recognized as a hallmark feature of aging, specific mechanisms by which aged T cells lose their capacity for effectively surveilling and navigating within complex microenvironments remain unclear^7–9^. To surveil, T cells require a set of core characteristics that include the capacity to: a) migrate, physically translating from one location to another; b) sense local signaling cues, which can be chemical cues such as cytokine gradients or physical cues such as local tissue stiffness; and c) deform, physically changing their morphologies or shapes as they polarize to move and squeeze around obstacles. Understanding whether aging impacts one or more of these core characteristics is critical to establishing a biophysical basis of T cell surveillance, and for developing effective strategies to modulate the surveillance behaviors of aged T cells.

T cells are inherently dynamic cell types. These highly migratory cells participate in continuous surveillance via the sampling of their local microenvironments to elicit specific T cell responses. Because immune responses require swift and timely reactions, T cell migration plays a critical role in how quickly a T cell finds its target and its capacity to follow local immune modulating signals, such as antigens and chemokine gradients^10^. Upon activation, T cells upregulate mechanisms and receptors, which modifies their migration behaviors. However, T cells must balance specific features of their activities, such as their migration speed or persistence and their ability to remain in a location for timely engagement with its target^11^. Because of this, T cells must adapt and tune their migration based on the spatial context of the microenvironment, resulting in a diverse repertoire of T cell migration behaviors. This range of T cell migration behaviors is generated through a combination of cell-intrinsic and cell-extrinsic factors, such as the rate of actin polymerization and the recognition of physical guidance cues (*e.g.,* ECM fibers, constrictions)^11–13^. However, it is unclear whether surveillance defects in aged T cells are primarily driven by a) dysfunctions in cell-intrinsic mechanisms that directly regulate their migration and deformability, b) progressive declines in their ability to sense and respond to cell-extrinsic factors, or c) a combination of both intrinsic and extrinsic factors.

Conventional approaches to profile the migration behaviors of T cells have relied heavily on aggregating data across multiple T cells to determine average values of migration parameters, for example, average speed, average persistence, and average turning angles. While computing average migration parameters across multiple T cells has provided critical insights into T cell migration, these approaches do not account for the range of heterogeneity in T cell migration. Recent advances combining single-cell imaging with machine learning now provide unprecedented tools for profiling migration behaviors of cells and simultaneously capturing the range of single-cell heterogeneities^14–18^. Furthermore, the capacity to profile T cell behaviors at single-cell resolution would enable deep characterization of multiple features of T cell surveillance simultaneously (*i.e.,* migration, deformability, and sensing in the same T cell). Given these possibilities, we sought to determine the effects of donor age on T cell surveillance behaviors.

In this study, we combined high-throughput time-lapse imaging with single-cell analyses to develop a suite of experimental and computational tools to profile the surveillance behaviors of T cells across a cohort of 51 healthy donors aged 18-93. Profiling the core characteristics of T cell surveillance, specifically, migration, deformability, and sensing, we found that T cells retained their capacity for spontaneous migration across the lifespan. However, aged T cells exhibited decreased cellular deformability and limitations in their ability to sense and respond to local signaling cues, such as secretions from neighboring cells. To evaluate whether aged T cells could be re-educated to exhibit ‘*young-like’* surveillance behaviors, we mechanically reprogrammed aged T cells using acute priming with elevated viscosity. Interestingly, aged T cells primed for seven days in extracellular fluid with an elevated viscosity exhibited ‘*young-like’* surveillance behaviors when seeded within a confined migration device and exposed to a stroma-derived factor 1 (SDF1) gradient. However, this ‘*young-like’* surveillance behavior was transient, lasting less than four days post-priming. While aged T cells exhibited enhanced surveillance behaviors, young T cells did not exhibit improvements in their behaviors, suggesting that they were already exhibiting optimal surveillance behaviors. Investigating why aged T cells exhibited enhanced surveillance behaviors, we found that acute priming at an elevated extracellular fluid viscosity initiated an outside-in cascade. This cascade was mechanistically linked to increased receptor expression and differential increases in membrane tension, cortical F-actin, and Actin-related protein 3 (Arp3) expression. These levels were comparable to levels observed in young T cells.

## RESULTS

### The capacity for T cells to migrate spontaneously is conserved during aging

Aging influences T-cell functions by impairing their ability to surveil effectively and navigate local microenvironments^6^. However, the phenotypic drivers governing these aging-related surveillance defects, as well as the mechanisms by which they develop in aging T cells, remain unclear. To develop a deep understanding of how the core characteristics of T cell surveillance change during aging, we first profiled the capacity of T cells to migrate spontaneously *ex vivo*. Prior studies have demonstrated that a variety of cell types in humans lose their capacity to migrate spontaneously with increasing age^19^. However, whether aging human T-cells also exhibit alterations in this ability to migrate, and whether these effects are extended to defects in sensing or deformation, remain poorly understood.

To investigate age-related changes in T cell migration, we isolated primary CD3^+^ T cells from whole blood of healthy human donors ranging in age from 18-93 years (**Fig. 1a**, **Supplementary Table 1**). We then profiled the spontaneous migration behaviors (*i.e.,* the capacity of cells to move) of individual T cells as a function of donor age. We denoted donors as young or old if they were 18-47 or 72-93 years old, respectively, and we refer to T cells from young donors as young T cells and T cells from old donors as aged T cells. To image and quantify single-cell migration behaviors, we immobilized T cells beneath a thin 0.5 mg/mL type I collagen hydrogel and acquired time-lapse tile scans (2×2) per position every 2 minutes, for a total of 4 hours. To quantify single-cell behaviors, we first segmented each cell using CellPose^47^, then tracked individual T cells for up to 4 hours per cell, after which we cut movies into unique 1-hour segments of continuous tracking, which we refer to as single-cell trajectories (see Methods) (**Fig. 1b**).

**Figure 1:**
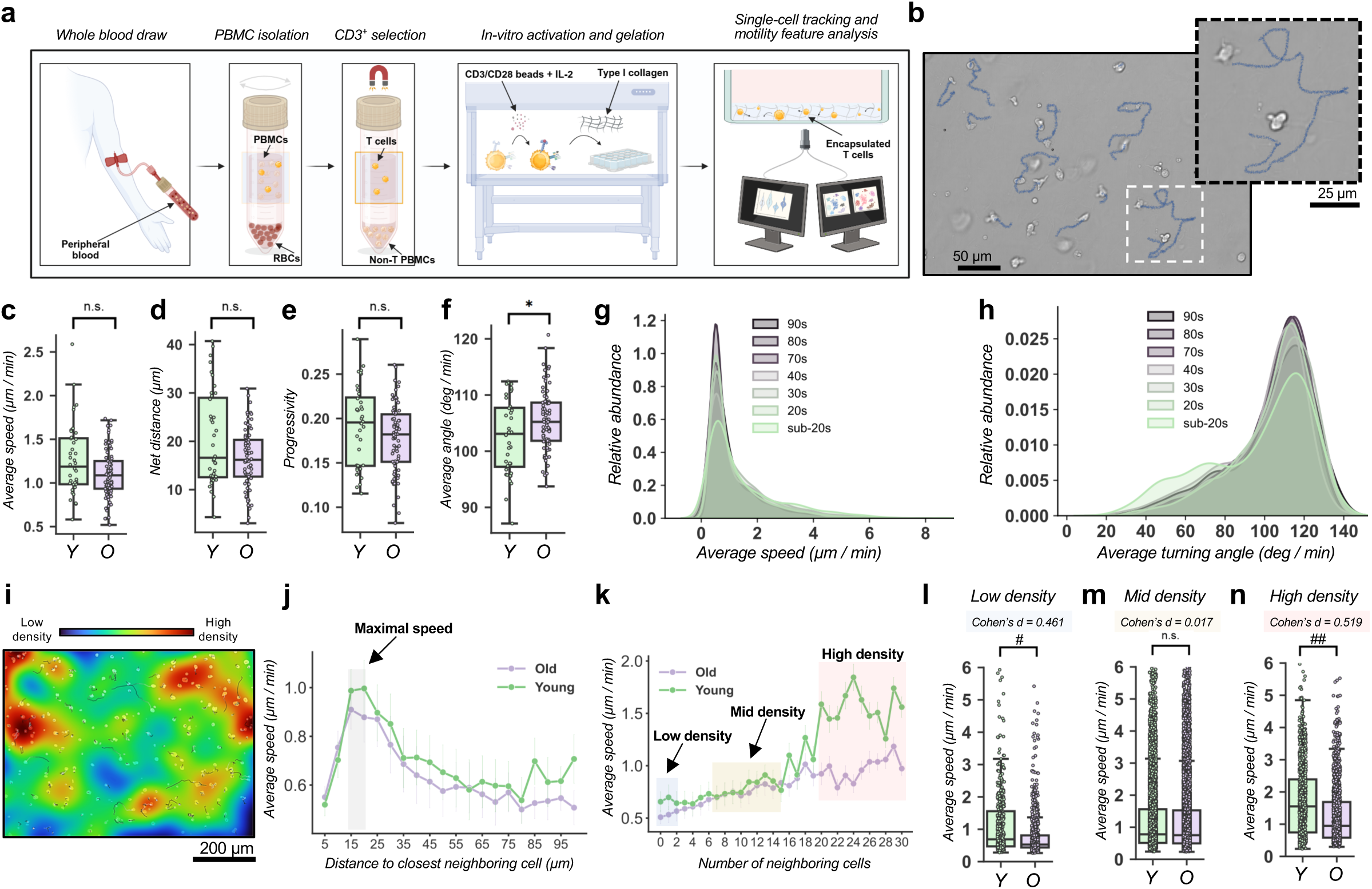
The capacity of T cells to migrate is conserved during aging. **a.** Schematic of experimental workflow to isolate, purify, culture, and measure the migration of human T cells derived from peripheral blood. **b.** Brightfield image of T cells migrating in 0.5 mg/mL type I collagen gels *ex vivo*. Blue lines indicate elapsed movement in a 1-hour trajectory. Inset follows one sample single-cell trajectory. **c.** Bulk analysis of select migration parameters, presented as one dot per technical replicate of young (*n* = 40) and old (*n* = 77) samples from 43 donors (15 young donors, 28 old donors): average speed, **d.** net distance, **e.** progressivity, and **f.** average turning angle. **g-h.** Kernel-density estimate plots of (*n* = 34,831) single-cell average speed (g) and average turning angle (h) binned by decades of donor age. Relative abundance refers to the density of occurrences at each motility feature value. **i.** Spatial cell density heatmap of local T-cell densities projected onto an image of migrating T cells. Blue denotes low density, and red denotes high density. **j.** Average speed of single cells (*n* = 34,831 cells) plotted as a function of the average distance to the nearest neighboring T cell. Grey shading refers to the distance range at which T cells reach maximal speed for both age groups. **k.** Average speed of single cells (*n* = 34,831 cells) plotted as a function of the average number of neighboring T cells. Blue, yellow, and red shading refer to regions of low, mid, and high cell density, respectively. **l.** Average speed of T cells at low spatial cell densities (2 or fewer neighbors; *n* = 1,155 cells), **m.** mid spatial cell densities (7-15 neighbors; *n* = 17,340 cells) and **n.** high spatial cell densities (20 or more neighbors; *n* = 2,334 cells). \**p* < 0.05; *n.s. Cohen’s d < 0.1;* #*0.2 ≤ Cohen’s d < 0.5; ##0.5 ≤ Cohen’s d < 0.8*.

Since T cell activation is known to influence migratory behaviors^11^, we evaluated whether *ex vivo* activation of T cells was necessary to thoroughly profile T cell migration. Upon activation using co-stimulation via CD3/CD28 and exogenous Interleukin-2 (IL-2; 50 ng/mL) for at least 3 days, we observed that non-activated T cells exhibited significantly reduced migration speeds, with a high fraction of non-migratory cells relative to activated cells (**Supplementary Fig. 1a-f**). Given this result, we performed all experiments using activated T cells. Across the forty-three donor samples used for migration experiments, we curated a total of 38,836 single-cell trajectories. For each of these trajectories, we computed thirty-three migration parameters, describing features such as the average speed, maximum speed, and average turning angle per cell (**Supplementary Table 2**).

After activating, imaging, and tracking all T cells for each donor, we pooled the data from all cells. We then performed a conventional analyss using average (bulk) migration parameters, comparing the two categories: young and old. For most measured migration parameters, we observed no significant differences between young or aged T cells. This finding suggests that aged T cells are capable of spontaneous migration comparable to that of young T cells (**Fig. 1c-e**). While the magnitude of T-cell migration (*i.e.,* how far they migrated or how fast) was conserved with age, the aged T cells tended to follow more tortuous paths, as indicated by a significant increase in average turning angle (**Fig. 1f**).

These results raised questions about whether subsets of T cells from young donors might exhibit differential migration characteristics, for example, do subpopulations of young T cells exhibit different migration speeds? This seemed possible because, although not statistically significant, we noted an apparent trend towards higher average migration speeds and net distance traveled for young T cells relative to aged T cells (**Fig. 1c, d**). To investigate this, we pooled all single-cell trajectories across all donors, then plotted the distributions of average speeds and average turning angles per decade (**Fig. 1g, h**). Results indicated that while the majority of T cells across the decades exhibited a similar mean value for average speed and average turning angle, there was a subpopulation of T cells in the younger decades, which exhibited higher average speeds and lower average turning angles, which we refer to as fast and persistent T cells.

In addition to quantifying the effects of age on the spontaneous migration of T cells, we considered whether biological sex or frailty status (based on the Fried frailty assessment for old donors^20^) differentially influenced T cell migration behaviors. Results indicated no significant differences in the migration behaviors of T cells from males versus females (**Supplementary Fig. 1g-l**), or from non-frail versus frail older adults (**Supplementary Fig. 1m-r**). Lastly, to assess whether the ratios of CD4+ to CD8+ T cells changed significantly as a function of age within our donor cohort, we performed flow cytometry experiments. Results indicated no statistically significant differences in the ratios of CD4+ to CD8+ T cells within the subset of donors evaluated (**Supplementary Fig. 2a-c**).

Together, these results indicate that the capacity for T cells to migrate spontaneously is conserved during aging. However, subpopulations of young T cells demonstrate enhanced migration behaviors

### Local cell density enhances T cell migration in an age-dependent manner

Given our observation that subpopulations of young T cells exhibit enhanced migration (**Fig. 1g-h**), we investigated factors potentially contributing to age-related shifts. Recent studies have shown that cell density can modulate the spontaneous migration of T cells^21,22^. For a comprehensive evaluation of the role of local cell density in T cell migration, we developed a robust single-cell approach. This assessment interrogates the spatial association of all cells within a given image field and quantifies, on a per-cell basis, using the location of every cell per time step and the number of neighboring cells. Then we reanalyzed the migration data to assess the effects of local cell density (see Methods). Despite seeding the same total number of cells (*n* = 15,000 cells/well), the spontaneous migration of T cells and their local cell-cell interactions (which can modulate their residence time in a location) influenced the formation of local cell neighborhoods. These high-density neighborhoods orchestrate signaling cascades via the secretion of signaling factors, *e.g.,* cytokines and chemokines, that further influence their migration^21,23^.

To visualize the formation of high-density neighborhoods, we performed spatial correlation analysis to dynamically map local cell densities of spontaneously migrating T cells across sequential frames per trajectory (**Fig. 1i, Supplementary Figure 3**). From this analysis, we quantified how the average speed in young and aged T cells changed as a function of distance to closest neighbor (**Fig. 1j**) and number of neighboring cells (**Fig. 1k**). Interestingly, both young and aged T cells achieved maximum average speeds when they were located 15-20 µm from their nearest neighbor, with a subsequent drop in average speed when they were further away (**Fig. 1j**). On average, across the 1-hour trajectories, T cells encountered up to thirty neighboring cells. Interestingly, the number of neighboring cells showed striking age-related differences (**Fig. 1k**). For instance, aged T cells migrating in either low (≤ 2 cell neighbors) or high (≥ 20 cell neighbors) density regions exhibited lower average speeds relative to young T cells; ∼20% and ∼50% reduction, respectively (**Fig. 1l, n**). However, aged T cells migrating within intermediate density regions (7 to 15 cell neighbors) exhibited comparable average speeds relative to young T cells (**Fig. 1m**).

These results highlight that, although young and aged T cells retain their migratory capacity, local cell density strongly influences T cell migration behaviors. Furthermore, the effect of local density has a greater effect on young T cells than aged T cells. This is indicated by the greater change in the magnitude of average speeds when young T cells are in high density regions relative to low density regions. These findings also suggest that the reduced effect of local density on aged T cells may be associated with a decreased capacity to sense and respond to local changes within their microenvironment.

### Aging induces a fractional redistribution across single-cell migration behaviors

In an earlier study, we developed a robust, data-driven framework named CaMI^14^ to robustly identify and classify single-cell migration behaviors and quantify the extent of migration heterogeneity across populations of cells. The data in Figure 1 (g-h) shows that T cells can adopt a wide variety of heterogeneous migration behaviors based on their biological context. To understand this heterogeneity, we performed a comprehensive single-cell analysis to characterize migration behaviors across young and aged T cells using a modified version of CaMI^14^ (**Fig. 2a**). Using CaMI allowed us to quantify the same thirty-three migration parameters described in Figure 1 (**Supplementary Table 2**) for each T cell within our dataset. We then pooled the data across all T cells (*n* = 34,831 cells), performed dimensionality reduction, and applied unsupervised k-means clustering to delineate groups of T cells with similar migration behaviors.

**Figure 2:**
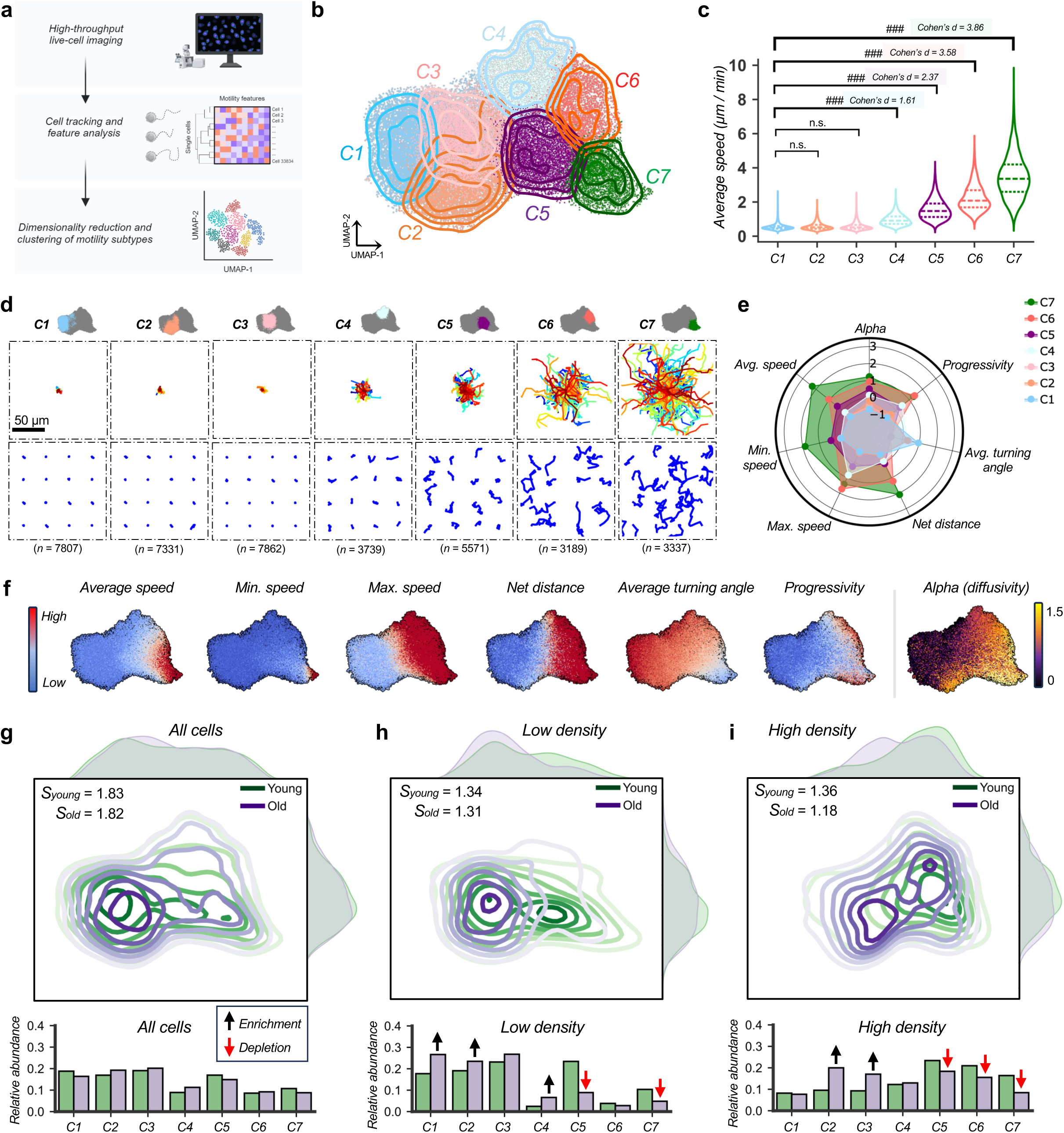
T-cell density induces a fractional redistribution among motility clusters in an age-dependent manner. **a.** Schematic depicting the workflow for single-cell dimensionality reduction of 33 motility features into a 2-dimensional space. **b.** UMAP and contour representation of (*n* = 38,836) single-cell motility patterns after unsupervised *k*-means clustering identifies seven distinct motility subtypes. Each dot represents one cell. **c.** Average migration speed of each *k*-means cluster. **d.** (*Top*) One-hundred representative trajectories of each cluster. (*Bottom*) Sixteen representative migration trajectories shown in a 4-by-4 grid for each cluster. The total number of cell trajectories per cluster is shown in parentheses. **e.** Radar plot illustrating differences in common migration parameters. **f.** Heatmap projection of select migration parameters onto the UMAP space. **g.** (*Top*) UMAP contour representation and (*bottom*) relative abundance of young and old donor-derived T-cell cluster subtypes. **h.** (*Top*) UMAP contour representation and (*bottom*) relative abundance of young and old donor-derived T-cell cluster subtypes when found at low densities (2 or fewer neighbors; *n* = 1,155 cells). **i.** (*Top*) UMAP contour representation and (*bottom*) relative abundance of young and old donor-derived T-cell cluster subtypes when found at high densities (20 or more neighbors; *n* = 2,334 cells). Permutation tests were run with 10,000 iterations. Black and red arrows denote relative enrichments and depletions, respectively. *S* denotes Shannon Entropy to calculate heterogeneity across multiple donors. *n.s. Cohen’s d < 0.1;* #*0.2 ≤ Cohen’s d < 0.5; ##0.5 ≤ Cohen’s d < 0.8; ###Cohen’s d ≥ 0.8*.

To reduce the dimensionality of our feature space, we performed Principal Component Analysis (PCA) across the thirty-three features, then extracted the principal components that explained 95% of the observed variance. These select principal components were used to create a two-dimensional, Uniform Manifold Approximation and Projection (UMAP) space, enabling us to visualize the complete ‘universe’ of T-cell migration behaviors in our sample. In parallel, we performed K-means clustering analysis on the same data, which yielded seven distinct T cell migration clusters: C1-7 (**Fig. 2b**). The optimal number of clusters was determined based on the inertia, silhouette, and the ratio of inertia over silhouette scores (**Supplementary Fig. 4a-c**). Although each migration cluster was defined based on the set of thirty-three measured features (**Supplementary Fig. 4d, e**), for ease of interpretation, we ordered the clusters based on the magnitude of their average speed. Thus, C1 identifies T cells with the lowest average speed while C7 identifies those with the highest average speed (**Fig. 2c**).

To visualize the T cell migration behaviors defined by each cluster, we plotted representative single-cell trajectories per cluster. This provided a qualitative view of cluster behaviors and highlighted the presence of distinct patterns of T cell migration (**Fig. 2d**). In addition to the representative trajectories, we plotted the magnitude of seven key migration parameters, further indicating the relative fundamental similarities and differences among the clusters (**Fig. 2e**). These seven key migration parameters included: 1) progressivity – the directional persistence of a cell, 2) average turning angle – the magnitude of the turning angle between subsequent steps, 3) net distance – the total distance a cell travels in 1 hour, 4) maximum speed – each cell’s maximum speed achieved within the 1-hour trajectory, 5) minimum speed – each cell’s minimum speed achieved within the 1-hour trajectory, 6) average speed – cells’ average speed achieved (distance/time) within the 1-hour trajectory, and 7) alpha – the slope of the mean-squared displacement indicating diffusive/random migration (alpha=1), sub-diffusive (alpha<1), or super-diffusive (alpha>1).

To visualize how these migration parameters varied, the UMAP shows the distribution of color-coded magnitudes of the seven migration parameters for each cell. Consistent with our expectation based on the migration behavior of C1 versus C7, cells on the lower right side of the UMAP denoted those with the highest average speed and lowest average turning angle (C7; **Fig. 2f**). In contrast, the far-left side of the UMAP denoted T cells with the slowest average speed and high average turning angles (C1). Notably, cells in C6 exhibited drastically lower average and minimum speeds relative to highly migratory T cells classified in C7. However, T cells classified in C6 were able to achieve significantly higher maximum speeds (**Fig. 2e, f**). Given this single-cell framework describing T cell migration, we revisited the data on the effect of activation on T cell migration. Consistent with our findings from the bulk migration analysis, non-activated T cells were not able to achieve the same extent of single-cell migration relative to activated T cells. Specifically, non-activated T cells were biased towards clusters showing low migration – C1, C2, C3, with low abundance in the mid and high migration clusters – C4, C5, C6, C7 (**Supplementary Fig. 1e, f**).

Given our previous finding that local cell density influences T cell migration behaviors (**Fig. 1i-n**), we questioned whether local cell density also skewed T cells towards specific migration clusters in an age-dependent manner. To evaluate this, we binned all T cells into three categories: all T cells (young and aged, independent of spatial density), T-cells in low density regions (≤ 2 cell neighbors), and T cells in high density regions (≥ 20 cell neighbors). Consistent with our findings in Figure 1, when local density is not considered both young and aged T cells displayed a similar likelihood of being classified in each of the seven migration clusters (**Fig. 2g**). However, young and aged T cells migrating in either low or high cell density regions exhibited striking differences in their migration behaviors with a fraction redistribution towards specific migration clusters (**Fig. 2h, i**). Aged T cells migrating in low density regions tended to be enriched in low and intermediate migration clusters C1, C2, C4, but were depleted in the high migration clusters C5 and C7 (**Fig. 2h**; *bottom)*. Furthermore, aged T cells migrating in high density regions were significantly enriched in C2 and C3 and depleted in C5, C6, and C7. Notably, migration clusters showing enrichment for aged T cells were depleted for young T cells, and *vice versa*. Interestingly, T cells migrating in low density regions exhibited reduced behavioral heterogeneity as determined by the Shannon entropy (*S_young_* = 1.34, *S_old_* = 1.31), relative to all cells independent of cell density (*S_young_* = 1.83, *S_old_* = 1.82) (**Fig. 2g-i**). T cells in high cell density regions also exhibited reduced behavioral heterogeneity. However, this was more pronounced for aged T cells relative to young T cells (*S_young_* = 1.36, *S_old_* = 1.18) (**Fig. 2i, Supplementary Fig. 5a-e**).

Collectively, these findings emphasize the differential effects of local cell densities on the elicited migration behaviors of young vs aged T cells. Furthermore, since T cells at high density are known to secrete higher levels of signaling factors relative to T cells in low density (*e.g.,* chemokines and cytokines)^21^, these results suggest a potential link between T cell migration, sensing, and response to microenvironmental signals.

### Aged T cells exhibit reduced morphological plasticity and deformability

To surveil effectively, T cells must be capable of dynamically deforming and changing their morphology to polarize and squeeze within constrictions and confined spaces^24^. For example, effector T cells disseminating from lymph nodes navigate complex networks of extracellular matrices^25^. Previous studies have shown that aging is associated with increased T cell stiffening, directly limiting their capacity to change their morphologies^26^. Thus, we considered whether aged human T cells within our study cohort may also exhibit reduced deformability. To assess this, we used two complementary approaches. First, we used an image-based approach to probe cell deformability based on spontaneous changes in T cell morphology observed between subsequent time steps, a measure we called morphological plasticity (see Methods). We assumed that, because the mechanical properties of T cells and their capacity to change morphology is intimately linked to the cytoskeleton (*i.e.,* actin)^27^, changes in T cell morphology can serve as a proxy for T cell deformability. We anticipated that this image-based approach would provide a high-throughput and scalable approach to measure T cell deformability. Second, using an orthogonal approach, we measured alterations in the shape of T cells using an optimized cellular microfluidic deformation assay in which deformability was defined as changes in the circularity of a cell when an external fluid force is applied^28^.

To profile the morphological plasticity of individual T-cells, we seeded single-cell suspensions in 0.5 mg/mL collagen hydrogels within a microfluidic device with a height of 10 µm (see Methods). Once cells were immobilized and attached, we acquired tile scans of young and aged T cells labeled with CellTracker (ThermoFisher Scientific). Images were acquired every 2 minutes for a total duration of 4 hours, out of which 1-hour segments were used for analysis. To determine the morphology of each T cell per frame, we performed segmentation using CellProfiler. Each generated T cell mask was tracked sequentially to establish a morphodynamic time series for each cell (see Methods). For each T cell, we assessed fourteen morphological features, including area, perimeter, eccentricity, and aspect ratio (**Supplementary Table 3**). Collectively, the fourteen morphological features defined changes in the size (*e.g.,* area, perimeter) and shape (*e.g.,* eccentricity, aspect ratio) of T cells. In an initial analysis, we profiled the average values of the morphology parameters comparing young versus aged T cells. We found that while the average values for most of the morphology parameters did not change markedly between young and aged T cells (**Supplementary Figure 6a-o**), the coefficient of variance for these parameters was significantly elevated in young T cells relative to aged T cells (**Supplementary Fig. p-ad**). The coefficient of variance denotes the rate of change in T cell morphologies across subsequent timesteps.

To quantify morphological plasticity based on the measured morphological parameters, we performed PCA on the fourteen morphological parameters for all T cells within our dataset (young and aged). From this, we generated a normalized PCA plot where a dot represents the morphology of each T cell per time step, then linking a cell’s subsequent dots created a morphodynamic time series for each cell (see Methods) (**Supplementary Fig. 7a, b**). Using this morphodynamic time series, we computed the morphodynamic plasticity as the speed of morphological changes per T cell within the PC space. A higher value of morphological plasticity indicated a greater change in morphology or a higher deformability. This composite measure also provided further robustness, as it was less sensitive to random fluctuations and noise in any given parameter.

Initially, we plotted representative morphologies of young and aged T cells every 10 minutes for 1 hour for cells that were navigating low, mid, and high density regions. Qualitatively, these data showed that aged T cells exhibited reduced morphological plasticity, especially when they were located within high density regions (**Fig. 3a**; *right*). Quantifying these results across all T cells, we found that when local T cell density was not considered, aged T cells showed significantly reduced morphological plasticity (**Fig. 3b**). Interestingly, in low density regions there was no difference in morphological plasticity between young and aged T cells (**Fig. 3c**), while aged T cells in mid and high density regions showed significantly lower morphological plasticity (*i.e.,* less deformability) than young T cells. Additionally, the greatest difference between young and aged T cells was observed in high density regions (**Fig. 3 d, e**). Together, these results suggest a relationship between a T cell’s ability to sense neighboring cells and its deformability (*i.e.,* its ability to regulate its morphology).

**Figure 3:**
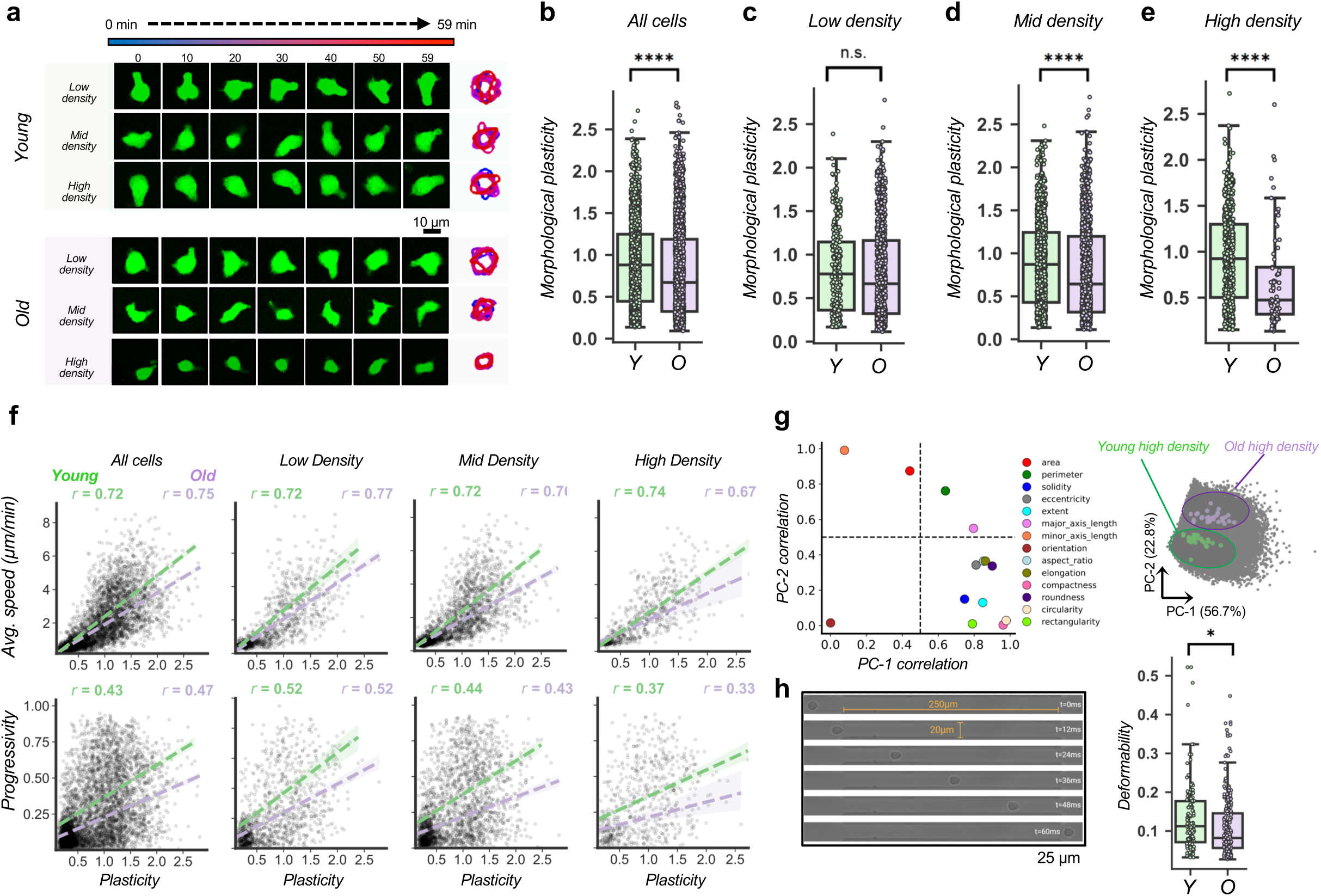
Aged T-cells exhibit reduced morphological plasticity and deformability. **a.** Representative images illustrating temporal changes in T-cell morphology. **b.** Quantification of age-dependent morphological plasticity, for **c.** T cells at low density (2 or fewer neighbors; *n* = 1,129 cells), **d.** for T cells at mid density (3 to 9 neighbors; *n* = 2,269 cells), and **e.** for T cells at high density (10 or more neighbors; *n* = 630 cells). **f.** Scatterplots and correlations of motility (average speed or progressivity) and morphological plasticity as a function of age and cell density. **g.** Correlations of principal components (PC-1 and PC-2) with individual features to indicate which features contribute to the variance in morphological plasticity. (*Right*) PC space of the young and old donor-derived cells at high density illustrated in (a). **h.** (*Left*) T cells are flowed through a microfluidic chamber and extrinsically deformed by fluid shear stress. (*Right*) Quantified extrinsic deformation of T-cells from young and old donors. \**p* < 0.05; \*\*\**p* < 0.001; \*\*\*\**p* < 0.0001.

Given the similarity in the effects of local density on the morphological plasticity and the migration of T cells, we computed correlations between morphodynamic plasticity and migratory parameters, specifically average speed and progressivity. We found that migration parameters such as average speed and progressivity were correlated with the morphodynamic plasticity in both young and aged T cells (**Fig. 3f**). The slope of fitted regressions was greater in young T cells compared to aged T cells. This indicated that young T cells could move faster than aged T cells when their morphological plasticity was matched. An alternative way to interpret this is that aged T cells needed to change their morphology to a greater extent than young T cells to migrate at comparable speeds. These morphological differences with age were conserved across all spatial densities (**Fig. 3f**).

To gain insight into specific morphological parameters that drive changes in T cell morphological plasticity, we computed correlations between PC1 and PC2 and each of the measured morphology parameters. Plotting the correlation of individual morphology parameters with PC1 (capturing 56.7% of the observed variance) and PC2 (capturing 22.8% of the observed variance) illustrated that a combination of size and shape changes were driving the observed changes in morphological plasticity (**Fig. 3g**; *left*). The results indicated that PC1 was highly correlated with many of the parameters describing changes in T cell shape such as circularity, roundness, and aspect ratio. On the other hand, PC2 was strongly correlated with parameters describing changes in T cell size such as the area, perimeter, and the major and minor axis lengths. Predictably, the morphological plasticity of young and aged T cells was defined by traversing distinct parts of the PC space (**Fig. 3g**; *left*). This differential localization of points belonging to the same T cells was greatest for young and aged T cells in high density regions. However, this difference diminished (more mixing of points in PC space) as the spatial density transitioned from high, to mid, to low density (**Supplementary Fig. 7a, b**).

Morphological plasticity is determined by intrinsically-driven control of cell surface velocity, cytoskeleton-mediated cellular mechanics, and water flow, all of which are closely coupled to actin cytoskeleton dynamics^29–31^. Since young T cells displayed elevated morphological plasticity relative to aged T cells, we sought to identify whether these cells had a greater capacity to be deformed by external forces. To assess this, we loaded suspended T cells into a custom, shear-based microfluidic device (**Supplementary Figure 7c, d**) and acquired high speed images of young and aged T cells as they experienced flow in defined geometries (**Fig. 3h**; *left*)^28^. The device was perfused at an optimized flow rate to deform the cells without causing them to rupture. This enabled us to detect significant changes in T cell morphologies before and after the application of fluid-induced deformation forces (**Fig. 3h**, *left*). Like our previous findings, we found that young T cells were more deformable compared to aged T cells (**Fig. 3h**; *right*). Furthermore, the agreement between these findings from both approaches provides validation that the changes in T cell moprhology (*i.e.,* morphological plasticity) can serve as an estimate of T cell deformability.

Collectively, these results indicate that aging reduces the intrinsic capacity of T cells to change their shape (morphological plasticity) as well as their capacity to be deformed by an external force. Furthermore, an inherent coupling exists between a T cell’s capacity to migrate and to deform, which influences its capacity to effectively surveil.

### Aged T cells exhibit defects in sensing, deformation, and migration when encountering chemical and physical gradients

The previous analyses indicate that aging reduces the ability of T cells to migrate and to deform as a function of their local environment. To develop an understanding of specific defects that may alter the capacity of T cells to sense signals within their local microenvironment, we exposed young and aged T cells to chemical and physical gradients within a microfluidic device (**Fig. 4a**). These devices contained a series of 3 µm-wide microchannels that connected the entry (seeding) channel to the exit (chemoattractant) channel^32^. This device enabled maintaining precise spatiotemporal control over the local environment of T-cells that are 5 to 10 µm in diameter. Additionally, passing through the 3 µm microchannels required T cells to squeeze as they moved within confinement, forcing them to change their cell morphology, cytoskeletal structure (*i.e.,* F-actin), and nuclear morphology during transit (**Fig. 4b**). Moreover, this configuration enabled direct testing of all three core characteristics needed for effective T cell surveillance, *ex vivo*. To transition from one side of the device to the other, a T cell must sense the gradient, deform to enter a channel, persistently migrate to the source of the gradient, exit the micro-channel, and re-adopt an unconfined morphology.

**Figure 4:**
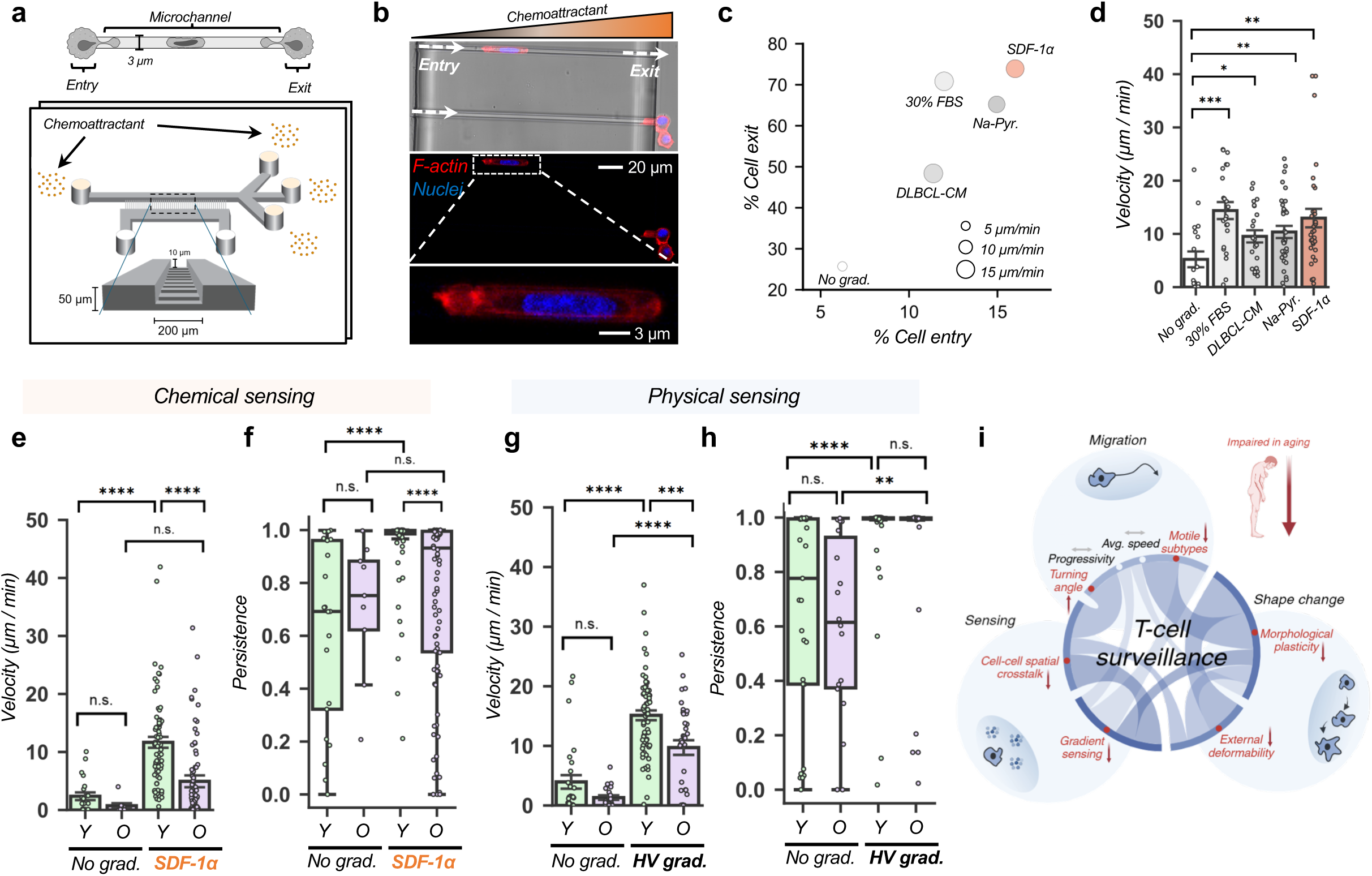
Aged T cells exhibit impaired chemotactic and physiochemical sensing. **a.** Schematic of engineered microchannels to assess T-cell migration, sensing, and deformation during confinement. **b.** Brightfield (*top*) and immunofluorescence (*middle*) images of T-cell chemotaxis towards a stromal derived factor-1α (SDF-1α) gradient. Inset (*bottom*) shows a T cell squeezing through a microchannel as it migrates towards an SDF-1α gradient. **c.** Scatter bubble plots of cell exit versus entry efficiency of T cells migrating under confinement without a gradient, or towards a gradient of high (30%) FBS, DLBCL-conditioned medium, sodium-pyruvate, or SDF-1α (calculated from *n* = 130 cells). **d.** Velocity of (*n* = 130) T cells migrating under confinement towards each type of chemotactic gradient. **e.** Velocity and **f.** persistence of young and old patient T cells migrating under confinement in the absence and presence of an SDF-1α gradient (*n* = 192 cells). **g.** Velocity and **h.** persistence of young and old patient T cells migrating in the presence of a control viscosity (CV; 0.77 cP) or high viscosity (HV; 8 cP) gradient (*n* = 152 cells). **i.** Graphical representation of how aging impacts T-cell surveillance. \**p* < 0.05, \*\**p* < 0.01, \*\*\**p* < 0.001, \*\*\*\**p* < 0.0001.

To identify an optimal T cell chemoattractant, we tested five candidate conditions: (1) no gradient, (2) high (30%) FBS, (3) B cell lymphoma-conditioned media (DLBCL-CM), (4) sodium pyruvate (Na-Pyr, 100μM), and (5) stromal-derived factor 1-alpha (SDF-1α, or CXCL12 (200 ng/mL)). For each condition, we quantified the entry-to-exit efficiency based on the fraction of cells entering and successfully exiting the microchannels (**Fig. 4c**). In addition to measuring entry-to-exit efficiency we computed the migration velocity of T cells transitioning from one side to the other within the microtubules (**Fig. 4d**). The results of these tests indicated that SDF-1α would be the best chemoattractant for evaluating the sensing ability of young and aged T cells. In a comparison of young and aged T cells responding to either no gradient or an SDF-1α gradient we found that in the absence of a gradient, both young and aged T cells exhibited similar velocities (∼2 µm/min) when migrating within the microchannels. However, in the presence of an SDF-1α gradient, young T cells exhibited a significantly higher velocity in migrating within the microchannels (∼10 µm/min on average) (**Fig. 4e**).

Although aged T cells also increased their migration velocity within the microchannels (∼4 µm/min), this change was not significant compared to their no-gradient controls. This chemotactic-driven increase in migration velocity of young T cells was particularly interesting considering that young T cells were ∼5-10% larger in size compared to aged T cells (**Supplementary Fig. 6a-c**, **Supplementary Fig. 6i**). Complementary measurement of migration persistence within the microchannels followed a similar trend indicating a strong increase in migration persistence in young T cells exposed to an SDF-1α gradient (**Fig. 4f**). Given the role of actin in regulating cell migration within confined channels^33^, we evaluated whether actomyosin contractility was essential for T cells to squeeze and migrate within the microchannels. We found that inhibition of actomyosin contractility via Rho-associated protein kinase (ROCK) with Y-27632 was sufficient to slow the SDF-1α-mediated migration from ∼10 µm/min to ∼2 µm/min in both young and aged T cells (**Supplementary Fig. 8a, b**).

To determine whether age-dependent changes were specific to chemical sensing, we repeated experiments using an elevated viscosity gradient to determine whether T cells could also sense changes in their physical environments. Standard cell culture media formulations have viscosities of approximately 0.77 cP. Using methylcellulose, we adjusted the viscosity of cell culture media, while maintaining other properties of the media (*e.g.,* osmolarity) (see Methods). Previous studies have shown that a variety of cell types (malignant and non-malignant) can respond to changes in viscosity^33^. To establish the viscosity gradient, we applied a high viscosity (HV) of 8 cP on one side (exit) and 0.77 cP on the other side (entry). The no-gradient control contained 0.77 cP on both sides. Interestingly, both young and aged T cells exhibited significant increases in their migration velocities and persistence when exposed to the viscosity gradient (**Fig. 4g, h**). However, young T cells achieved a significantly higher migration velocity relative to aged T cells with no changes in their persistence.

Collectively, these results suggest that the capacity to sense changes in the local environment is a potential limiting characteristic for effective T cell surveillance. Furthermore, the decreased capacity of aged T cells to sense signals within their microenvironment is intimately coupled with their migration and deformation capabilities, as indicated by our assessment of local spatial density effects (**Fig. 4i**).

### Viscous priming transiently reprograms surveillance behaviors in aged T cells

Prolonged incubation and priming with elevated viscosity modulate the migration and behaviors of cells^31,33,34^. Furthermore, given the strong response to a viscosity gradient by aged T cells, we evaluated whether priming with elevated viscosity could induce ‘young-like’ surveillance behaviors in aged T cells. Recent work has shown that multiple cell types are mechano-sensitive and able to exhibit vastly altered cytoskeletal structures and water fluxes, leading to increased migration capability^35^. Recently, viscous priming has been established as an effective tool to drive changes in cellular migration, morphology, and cell fate through modulation of actomyosin activity, or transmembrane channels, depending on the cell type^31,34^. Whether this approach could be applied to mechanically reprogram aged T cells remains unknown.

To address this, we cultured activated young and aged T cells in 0.77 cP or 8 cP, respectively (**Fig. 5a**). After 7 days, we thoroughly washed the cells using PBS, and profiled their surveillance behaviors (sensing, deformation, and migration) within confined microchannels, as previously described (**Fig. 4a**). The results indicated strikingly that aged T cells primed with elevated viscosity exhibited migration velocities and persistence comparable to young T cells (**Fig. 5b, c**). Specifically, aged T cells primed with elevated viscosity (8 cP) migrated towards the SDF-1α gradient at ‘*young-like’* velocities (∼10 µm/min) and persistence (∼1). Interestingly, young T cells primed with either 0.77 cP or 8 cP exhibited unchanged migration velocity and persistence, suggesting that young T cells may already function at near optimal levels. Given these results, we assessed how long this mechanical reprogramming persisted. To evaluate this, we primed young and aged T cells for seven days and, after washing with PBS, cultured the cells in regular 0.77 cP media for 1, 4, or 7 days. Interestingly, aged T cells retained their ‘*young-like*’ surveillance behaviors with elevated migration velocity and persistence at one day post-priming (**Fig. 5d, g**). However, the mechanically-induced reprogramming did not persist to four (**Fig. 5e, h**) or seven days post-priming (**Fig. 5f, i**).

**Figure 5:**
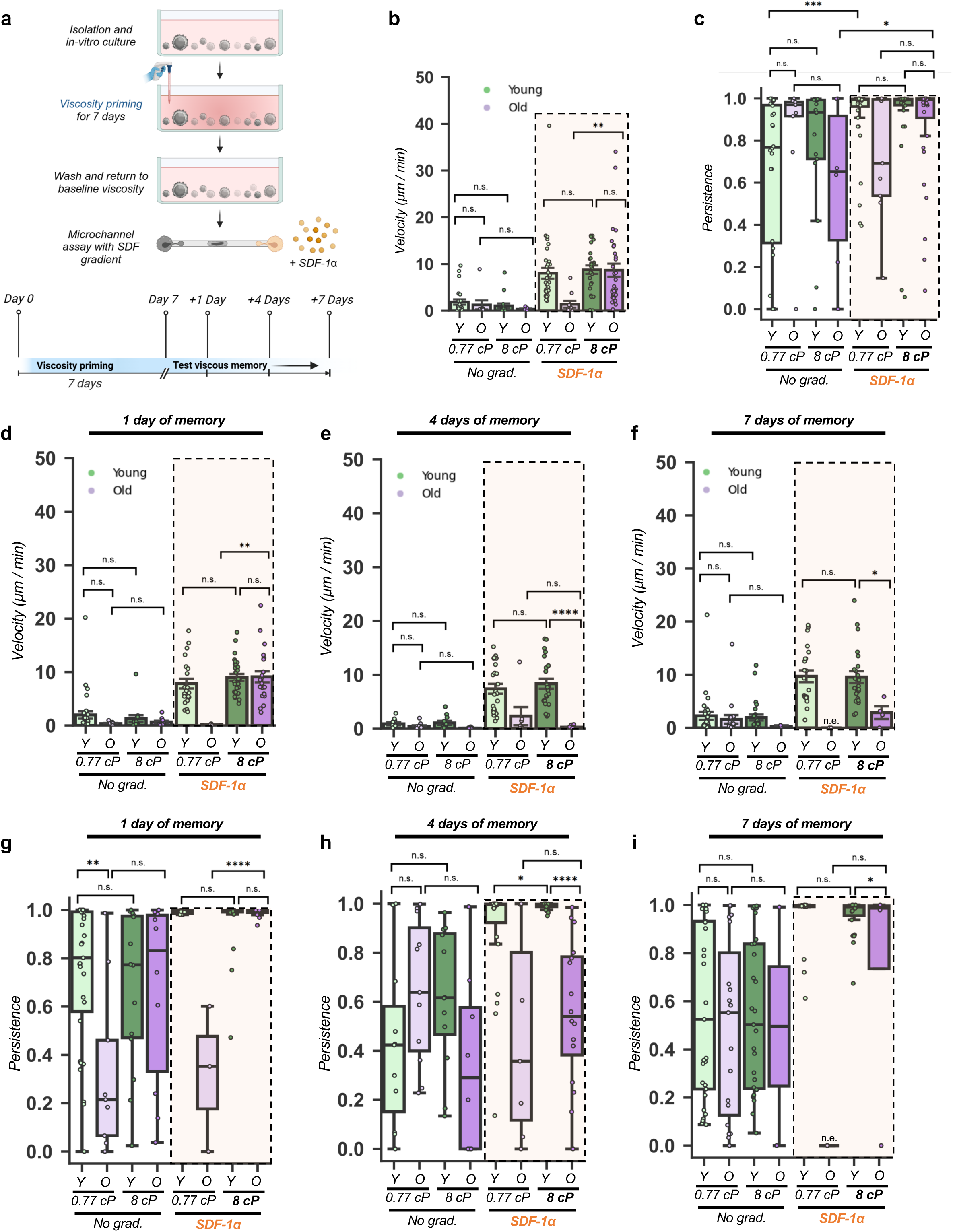
Viscous priming transiently rescues *young-like* surveillance behaviors in aged T cells. **a.** Schematic to precondition T cells in control or high viscosity for 7 days and test viscous memory after an additional 1, 4, and 7 days. **b.** Velocity and **c.** persistence of control viscosity (0.77 cP)- and high viscosity (8 cP)-primed young and aged T cells migrating under confinement in the absence and presence of an SDF-1α gradient (*n* = 160 cells). **d.** Velocity of young and aged T cells migrating under confinement in the absence and presence of an SDF-1α gradient after 1 day (*n* = 139 cells), **e.** 4 days (*n* = 111 cells), and **f.** 7 days (*n* = 128 cells) since priming in control (0.77 cP) or high viscosity (8 cP) media. **g.** Persistence of young and old donor-derived T cells migrating under confinement in the absence and presence of an SDF-1α gradient after 1 day, **h.** 4 days, and **i.** 7 days since priming in control (0.77 cP) or high viscosity (8 cP) media. \**p* < 0.05; \*\**p* < 0.01; \*\*\**p* < 0.001; \*\*\*\**p* < 0.0001.

These findings suggests that priming aged T cells with elevated viscosity induces a transient mechanical reprogramming to ‘*young-like*’ levels, providing evidence for mechanical modulation of T cell behaviors.

### Viscous priming modulates aged T cell behaviors through an F-actin—Arp3 axis involving elevated receptor expression and increased membrane tension

We next sought to understand the biological mechanism driving the mechanical reprogramming of aged T cells. As we and others have demonstrated, T cells can respond to SDF-1α, a common chemokine involved in immuno-stromal crosstalk. Furthermore, SDF-1α exposure is known to drive increased expression of genes related to T cell adhesion, changes in T cell morphology, and chemotaxis^36,37^. Interestingly, priming T cells in 8 cP media increased the abundance of CXCR4, a G-protein-coupled chemokine receptor of SDF-1α on the surface of T cells. However, no differences were exhibited in expression in primed young or aged T cells (**Fig. 6a**).

**Figure 6:**
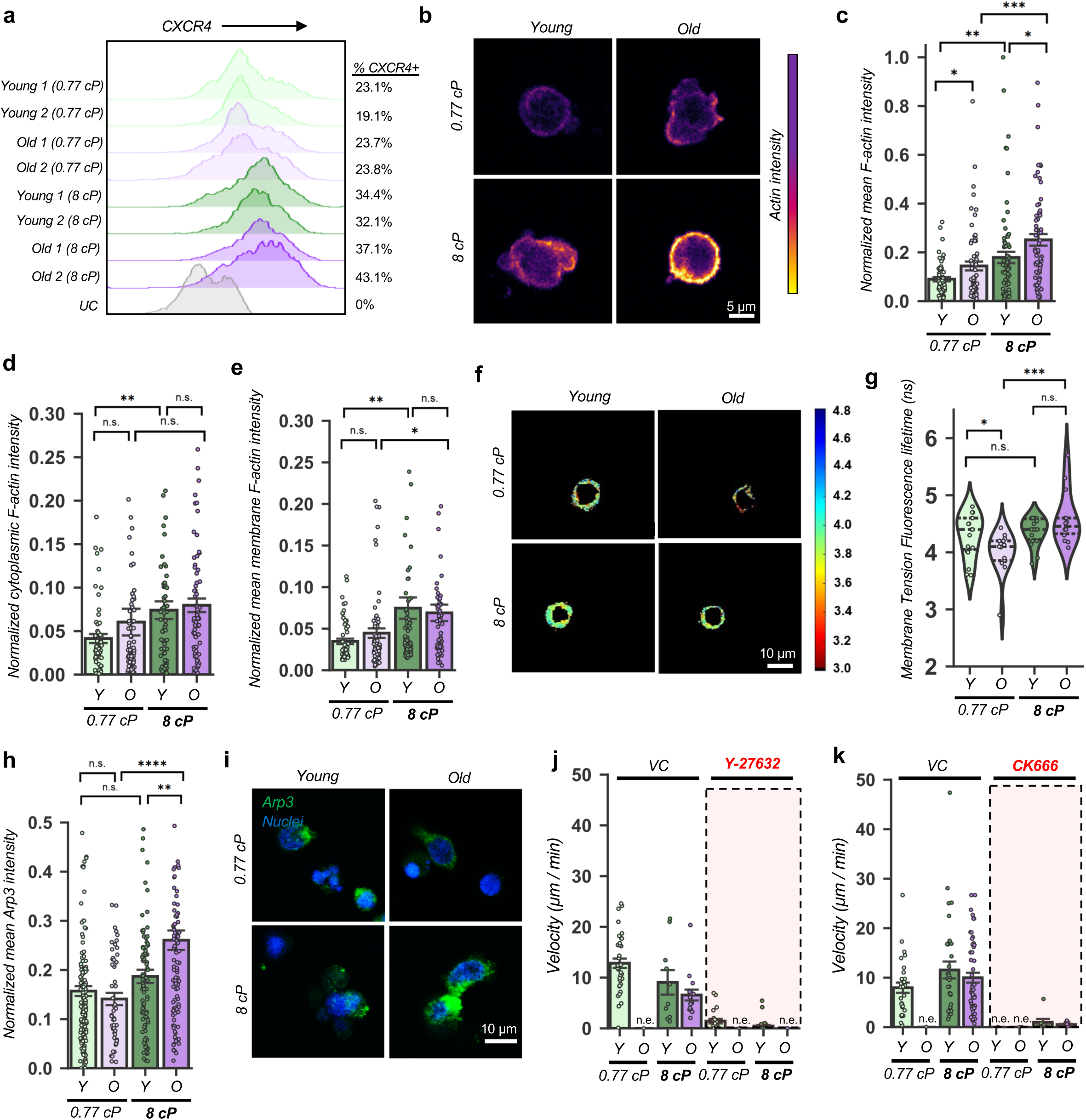
Viscous priming enhances sensing in aged T cells. **a.** Flow cytometry quantification of the SDF-1α receptor (CXCL4) in young and old donor-derived T cells after 7 days priming with control (0.77 cP) or high (8 cP) viscosity (*n* = 1,138,445 cells). Two donors of each age group were assessed. *UC*, unstained control. **b.** Representative immunofluorescence images of F-actin content in young and old donor-derived T cells after 7 days of priming with 0.77 cP or 8 cP viscosity. **c.** Quantification of mean F-actin content present in cells, **d.** specifically in the cytoplasm, and **e.** at the cell membrane of young and old donor-derived T cells after 7 days of priming with 0.77 cP or 8 cP viscosity (*n* = 263 cells). **f.** Representative images and **g.** quantification of cellular membrane tension in young and old donor-derived T cells after 7 days of priming with 0.77 cP or 8 cP viscosity (*n* = 66 cells). **h.** Quantification of mean Arp3 expression (with **i.** representative images) in young and old donor-derived T cells after 7 days of priming with 0.77 cP or 8 cP viscosity (*n* = 408 cells). **j.** Velocity of control viscosity (0.77 cP)- and high viscosity (8 cP)-primed young and old donor-derived T cells migrating under confinement in response to an SDF-1α gradient during treatment with either Y-27632 or **k.** CK666 compared to vehicle control (*n* = 145 cells). n.e., no entry of cells. \**p* < 0.05; \*\**p* < 0.01; \*\*\**p* < 0.001; \*\*\*\**p* < 0.0001.

Binding of SDF-1α to CXCR4 is known to induce actin-associated cytoskeletal remodeling^38^. Previous work demonstrated that elevated viscosity drives tighter macromolecular crowding, which enhances actin polymerization and causes cells to form denser, reinforced actin networks to counteract viscous forces^33,39^. Quantifying F-acting content in primed T cells revealed a significant increase in both young and aged T cells (**Fig. 6c**, **Supplementary Fig. 8c**). Although this increased F-actin was observed both in the cytoplasm and at the cortex (*i.e.,* membrane related), we observed a greater increase in cortical F-actin at the cell membrane (**Fig. 6d, e**). Live cell measurement of membrane tension further reinforced this finding, as viscous priming in 8 cP recovered membrane tension of aged T cells to ‘*young-like*’ levels. However, the membrane tension in young T cells remained unchanged (**Fig 6f, g**)

Intracellular Arp2/3 complex is closely linked with changes in membrane tension, as it restructures the polarization of intracellular actin filaments to generate force^40^. As such, the Arp2/3 complex is postulated to drive chemokine-driven T cell migration and morphological changes via modulation of actin polymerization^41^. Given the observed age-dependent increase in membrane tension in tandem with elevated F-actin, we investigated whether Arp2/3 also played a role in the mechanical reprogramming of aged T cells exposed to elevated viscosity. Qualifying the expression of the Arp3 subunit, we observed a differential increase in Arp3 expression in aged T cells primed with elevated viscosity (**Fig. h, i**). Furthermore, the downregulation of Arp3 is shown to reduce cell migration, actin remodeling, and lamellipodia formation, thereby defining an F-actin—Arp3 axis, which influences immune dysfunction and potentially contributes to the observed age-related defects in T cell surveillance^42,43^. To evaluate this notion, we validated the importance of the F-actin—Arp3 axis by inhibiting actin contractility and Arp3 activity in young and aged T cells primed with elevated viscosity. The results showed that inhibition of actomyosin contractility with ROCK inhibitor Y-27632, and Arp3 expression with CK-666 was sufficient to completely attenuate the ‘*young-like*’ surveillance behaviors in primed aged T cells to baseline levels (**Fig. 6j, k, Supplementary Fig. 8d, e**).

Altogether, these results highlight the role of the F-actin—Arp3 axis in regulating surveillance behaviors in aged T cells.

## DISCUSSION

Aging is a complex, multifaceted process accompanied by abundant changes at the molecular and cellular scales^44^. Conventional measurements of cell migration, which tend to rely on bulk analyses of averaged parameters are often limited in capturing subsets of cells that share migration properties across biological contexts. Consistently, our conventional (bulk) analysis of T-cell migration did not reveal strong age-related differences between young and aged T-cells, highlighting the importance of single-cell approaches (**Fig. 1, 2)**. Furthermore, our spatial density analysis was able to differentiate shifts in migration behaviors between young and aged T cells. These sensing-associated differences were also linked to morphological plasticity, which revealed that aging reduces the capacity of T cells to regulate their morphologies (**Fig. 3**). We determined that this dysregulation of sensing in aged T cells was not specific to a single signal (**Fig. 4**), and defective sensing and migration in aged T cells could be reprogrammed using elevated viscosity (**Fig. 5, 6**). In summary, priming aged T cells with elevated viscosity increases chemokine receptor presentation, increases F-actin content, boosts Arp3 expression, and restores membrane tension, inducing ‘*young-like*’ surveillance behaviors. In summary, our results present a framework to explain age-related defects in the surveillance behaviors of aged T cells, with a key limitation of T cell function in older adults being the age-related loss of T cell sensing.

Together, this work also provides insights and considerations for engineering behaviors in aged T cells for applications in immunotherapies. These findings suggest that unlike young T cells, aged T cells are significantly limited in their capacity to deform and sense signaling cues within local microenvironments. In addition to engineering effective chimeric antigen receptors (CARs), for instance, in various cancers, considerations should be made to engineer enhanced migration, deformability, and sensing traits in CAR T cells from older donors. These can be achieved in a variety of ways, for instance, enhancing CAR T cell migration could be achieved by including velocity receptors^45^. However, developing approaches to specifically enhance deformability and sensing require additional studies. While we have shown that mechanical reprogramming with elevated extracellular viscosity as a putative solution, the transient nature of the enhanced surveillance behaviors may be insufficient to see a sustained effect. As a result, additional studies are needed to develop approaches to sustain these enhanced behaviors.

While this work highlights key findings, additional studies are needed to further understand the downstream effects of viscous priming on the transcriptional and epigenomic states of aged T cells, and whether additional strategies can be applied to prolong the mechanical reprogramming. Strategies could include using small molecule drugs or genetic engineering tools to target genes or epigenetic modifications that are induced by viscous priming. This can be achieved by having a deeper understanding of the molecular changes as a function of priming kinetics (*i.e.,* baseline, primed state, post-prime). Critical questions could include: a) are there transcriptional or epigenetic changes in T cells between baseline (activated) and post-priming, and can these be modulated? b) does priming T cells with elevated viscosity for longer durations or cycles viscosity increase the memory of enhanced behaviors in aged T cells? and c) does the priming of T cells with elevated viscosity in aged T cells modulate activation and T cell function (e.g., cell killing)? Furthermore, this study utilized pan T cells. However, gaining deeper insights into whether subsets of aged T cells, (*i.e.,* CD4^+^ vs CD8^+^) exhibit differential responses to priming could be critical for developing next-generation T cell therapies. Finally, further research is required to determine whether age-related defects in T cell surveillance behaviors are extended to other immune cell types such as B cells, monocytes, macrophages, or neutrophils.

## MATERIALS AND METHODS

### Cell culture and hydrogel fabrication

Human peripheral blood mononuclear cells (PBMCs) were isolated from peripheral blood of 51 de-identified donors of age 18 to 93 (**Supplementary Table 1**) using density-dependent centrifugation (**Fig. 1a**)^46^. CD3^+^ T cells were purified from the PBMC population using a magnetic bead separation kit (Invitrogen), resuspended in complete media that contained RPMI, 10% FBS, 1% pen/strep, 10 mM HEPES, 1 mM sodium pyruvate (all from Gibco), and 4 ng/mL 2-mercaptoethanol (Sigma-Aldrich). Activated T cells were prepared using Activation Media, which was composed of complete media, ImmunoCult Human CD3/CD28 T Cell Activator (25 µL/mL; STEMCELL Technologies), and human recombinant IL-2 (50 ng/mL or 500 Units/mL; STEMCELL Technologies) at a density of 1×10^6^ cells/mL. T cells were activated for at least three days prior to use in experiments. Media of prescribed control (0.77 cP) and elevated (8 cP) viscosities were prepared by dissolving the appropriate ratio of a 3% methylcellulose solution (R&D systems) in IMDM (Life Technologies) that contained 10% FBS and 1% pen/strep before adding 25 µL/mL ImmunoCult Human CD3/CD28 T Cell Activator and 50 ng/mL IL-2.

To form hydrogels, rat tail type I collagen (Corning) was neutralized to pH 7-7.5 using 0.2 M NaOH and diluted to a final concentration of either 0.5, 1, or 4 mg/mL using a combination of 10× HBSS (Sigma-Aldrich), water, and complete RPMI. T cells were centrifuged and resuspended in collagen gels at a density of 150k cells/mL, plated in 96-well plates (100 µL/well), allowed to settle at 4°C for 20-30 min, polymerized at 37°C for 1 hr., and rehydrated with Activation Media.

### Live cell microscopy and image processing

Brightfield and fluorescence live-cell imaging for T-cell motility and morphological plasticity was performed using a Leica Stellaris 5 confocal microscope equipped with a 20×/0.75 NA objective and a humidified chamber (Tokai Hit, 37°C and 5% CO_2_). Images of T cell-laden gels were acquired every 2 min for a total of 4 hr. at 1024×1024 scanning resolution. Cell boundaries were segmented, and cell features were quantified in CellPose and CellProfiler^47,48^. Cell centroid movements were tracked using Trackpy, a Python package based on the Crocker-Grier algorithm, to classify motile trajectories for 30 successive frames (i.e., 1 hour) and quantify 33 single-cell motility features (**Supplementary Table 2**).

### Spatial analysis of T-cell neighborhoods

To compute local cell neighborhoods, we defined a 200 pixel (∼113 μm) search radius around each single cell and counted the number of unique neighboring cells that entered this search radius within each 1 hour single-cell trajectory. Quantification of cell-to-cell proximity on a continuous scale was performed by constructing centroid distance matrices between the cell of interest and each unique neighboring cell per frame. The final time series data computed the distance between the cell of interest and each of the identified neighboring cells, and we calculated the average shortest distance to the first neighboring cell and the average total number of neighboring cells for each single-cell trajectory.

### Dimensionality reduction and clustering

To prepare data for dimensionality reduction, all motility features were standard-scaled (z-scored) and log-normalized, and PCA (Principal Component Analysis) was used to capture the minimum number of principal components that explained 95% of the variance in the dataset. Uniform Manifold Approximation and Projection (UMAP) was used to condense the 33 motility features into a 2-dimensional plane in the Euclidean coordinate system, in which each dot on the UMAP space represents one single-cell trajectory. Similar behaviors of T-cell trajectories were grouped together using an unsupervised *K*-means clustering algorithm. A total of seven unique behavioral clusters were identified based on inertia, silhouette, and inertia/silhouette plots (**Supplementary Figure 4a-c**). Clusters were named in order of increasing average speed (**Figure 2c-d**).

### Randomized permutation for cluster enrichment and depletion

To quantify *K*-means statistically-significant cluster enrichment and depletion, we applied a randomized permutation test for each age (young and old) and cluster (*C1*-*C7*) by calculating the fraction of abundance as an observed statistic, *t_obs_*. At each iteration of the test, random shuffling of the *K*-means cluster column within the entire dataset was performed while fixing the biological (age) group column, and we recomputed the fraction of abundance for each group and for each cluster as a simulated statistic, *t_sim_*. The *p*-values for both enrichment and depletion are computed as follows:

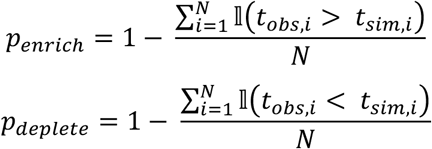

Here, *p_enrich_*, and *p_deplete_* are the *p*-values for enrichment and depletion, respectively, *N* signifies the total number of iterations (*N* = 1,000), and 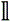 is the indicator function. The indicator function is equal to 1 if the condition is true, and equal to 0 if the condition is false.

### Microfluidic device fabrication for measurement of morphological plasticity

Silicon molds were fabricated using standard photolithography procedures. Photomask designs were drafted in AutoCAD and ordered from HTA Photomask. Molds were fabricated according to the manufacturer’s instructions for the SU-8 3010 negative photoresist. Briefly, two layers of photoresist were spin-coated on a silicon wafer (IWS) at 500 rpm for 7 s with an acceleration of 100 rpm/s, and at 2,000 rpm for 30 s with an acceleration of 300 rpm/s, respectively. After a soft bake of 4 min at 95°C, the desired microfluidic pattern was etched using UV light to yield feature heights of 10 µm.

A 10:1 ratio of polydimethylsiloxane (PDMS; Dow Sylgard) and curing agent was mixed thoroughly, poured onto each silicon wafer, and cured at 80°C for at least 45 min. Devices were sonicated in 100% isopropyl alcohol (IPA) for 5 min and dried using a compressed air gun. Microfluidic devices were bonded to 6-well glass bottom plates (#1.5H Glass; Cellvis) using oxygen plasma exposure for 70 s, then baked at 80°C for an additional 45 min to ensure complete bonding.

### Measurement of morphological plasticity

To quantify cellular morphodynamics, we constructed a two-dimensional morphology space using principal component analysis (PCA) on all multi-dimensional morphological features (**Supplementary Table 3**). This approach extracted the first two principal components to maximize (95%) variance in the dataset, while minimizing covariance. As a visual, we mapped cellular trajectories at each frame throughout this morphology space and calculated morphological plasticity as the mean instantaneous speed along the trajectories as a cumulative quantitative measure of morphological change over time.

### Microfluidic device to measure external deformability

The fabrication, preparation, and analysis of the MCDS procedures were conducted as previously described^28^. In short, PDMS devices, cast from a positive mold wafer, were permanently adhered to a glass cover slip and bonded overnight at 80°C. The following day, PDMS devices were plasma treated, prepared with 8% Pluronic F68 (P1300, Sigma-Aldrich), and rinsed with Milli-Q water. Suspended cells were centrifuged and resuspended in deformation fluid comprised of 0.5% methylcellulose (Sigma-Aldrich), 0.5% BSA (Sigma-Aldrich), and 0.1% sodium azide (Sigma-Aldrich). The suspension was transferred to a syringe (VWR) and mounted in a pump. With the PDMS device mounted on the microscope stage with the cell suspension and deformation fluid tubing attached, flow was initiated with a 3.5 min delay to allow the system to equilibrate. Following equilibration, cell deformation within the deforming channel of the MCDS was captured using a highspeed camera (MIRO-C210; Phantom). Acquired files were then exported, and cell deformation was quantified via tracing in FIJI/ImageJ.

### Microfluidic device fabrication for T-cell chemotaxis during confinement

Microfluidic devices were designed, fabricated, and bonded as previously described^33^. In summary, photolithography was used to pattern microchannels onto a silicon wafer. The microfluidic devices contained microchannels of prescribed length (200 µm), height (10 µm) and width (3 µm). PDMS was poured onto the silicon wafer and polymerized for ∼75 min at 85°C. A 5 mm-diameter biopsy punch was used to create 6 wells for cell seeding and media changes in later steps. PDMS devices were sonicated in ethanol for 5 min, washed with DI water, and activated with a plasma cleaner (Harrick). Devices were activated with a plasma cleaner and subsequently bonded to 25 x 75mm glass slides (Electron Microscopy Sciences) that were washed previously with ethanol and DI water. The glass slides were coated with 20 μg/mL type-I rat tail collagen (Gibco) for 1-1.5 hr. at 37°C prior to cell seeding.

In selected assays, 20,000 cells (cell density: 1,000 cells/µL) were added to the seeding well of the inlet channel in the microfluidic device (3×10 µm), creating pressure-driven flow of cells across the length of the seeding region. As the cells reached the outlet channel, an aliquot comprising 6.67 µL of cell suspension was moved to the outlet to balance the flow rate. Devices were incubated for ∼10 min to allow cells to adhere at the entrances of microchannels. After cells adhered, 60 µL of SDF-1α-containing media (200 ng/mL) was added to each of the top 4 wells of the device (**Fig. 4e**), and control media was added to the bottom 2 wells of the device to generate an SDF-1 α gradient. To create a physicochemical gradient, this process was repeated by using chemokine-free media of 0.77 cP viscosity in the downstream wells, and chemokine-free media of 8 cP viscosity upstream (**Fig. 4f**). Cells in microchannel confinement were imaged and tracked every 2 min for up to 2 hr. on an inverted Nikon Eclipse Ti microscope (Nikon, Tokyo, Japan) with automated controls (NIS-Elements; Nikon) and a 10×/0.45 NA Ph1 objective. A temperature- and CO2-controlled stage-top incubator (Tokai Hit) was utilized to maintain cells at 37°C with 5% CO2 during experiments. Cells were manually tracked in ImageJ using the Manual Tracking plugin (National Institutes of Health) and analyzed in MATLAB (v. R2022b; Mathworks).

### Immunofluorescence staining

For immunofluorescence staining, T cells were affixed to the bottom of 96-well-plates using Cell-Tak Cell and Tissue Adhesive (Corning), immobilized in 4 mg/mL collagen gels, or retained in microchannel confinement devices. All cells were fixed with 4% formaldehyde (ThermoFisher Scientific) for 15 min, washed three times with PBS, then permeabilized for 1 hr using blocking buffer, which was composed of 0.1% Triton X-100 (Millipore Sigma) and 2% BSA (Sigma) in PBS. Primary antibody solutions were prepared in blocking buffer before overnight incubation at 4°C: Phalloidin-647 (1:2000 ratio, Invitrogen), Arp3 (1:200 ratio, Abcam), or Hoechst (1:2000, Invitrogen). The next day, T cells were washed with PBS three times, incubated for 1 hr. with Alexa Fluor 488 goat anti-mouse immunoglobulin G (IgG) H+L antibody (1:200 ratio, Invitrogen), washed twice with PBS, then imaged on a Nikon Eclipse Ti equipped with a 10×/0.45 NA Ph1 objective or a Leica Stellaris 5 confocal microscope equipped with a 20×/0.75 NA objective.

### Flow cytometry

Surface antigen staining of CD4, CD8, or CXCR4 was performed on young and old donor-derived T cells. All cells were prepared in the same manner for staining and fluorescence acquisition, differing only in the specific antibody panel. For the CD4/CD8 T-cell subtype quantification panel, the following antibodies were used: anti-human CD3-FITC (Biolegend), anti-human CD4-PE (Biolegend), and anti-human CD8a-APC (Biolegend). For the CXCR4 quantification panel, CD184 (CXCR4) monoclonal antibody APC (eBioscience) was used. Prior to staining, cells were removed from culture and washed twice with PBS. Cells were then incubated for 30 mins with a flow cytometry blocking buffer (3% BSA in PBS) to reduce nonspecific antibody binding. For surface antibody staining, cells were resuspended in FACS buffer (1% BSA, 0.09% sodium azide) and incubated with fluorochrome-labeled antibodies targeting surface antigens for 1 hr. at 4°C while protected from light.Appropriate compensation controls were calibrated for each antibody using RayBright Compensation beads. Following antibody conjugation, cells were washed twice with fresh PBS. All dead cells were excluded from the analysis using propidium iodide. Fluorescence readouts were acquired on a FACS Canto Flow Cytometer (BS Biosciences) and analyzed using FlowJo version 10.9.0 (FlowJo LLC).

### Membrane tension quantification

T cells were immobilized in rat-tail collagen type I gels of high density (4 mg/mL) and stained with the live-cell membrane tension probe Flipper-TR (Spirochrome, 0.5 mM) and imaged immediately. Confocal fluorescence lifetime imaging microscopy (FLIM) of cells stained with Flipper-TR (0.5 mM, Spirochrome) was conducted by following previously established protocols using a Zeiss LSM 780 microscope coupled with a PicoQuant system^34^. This system included a PicoHarp 300 time-correlated single-photon counting (TCSPC) module, two hybrid PMA-04 detectors, and a Sepia II laser control module. FLIM data were processed and analyzed as detailed in previous reports.

### Preparation of microchannel inhibitors

Stock solutions of Y-27632 and CK666 were reconstituted by dissolving each compound in DMSO to concentrations of 10 mM and 100 mM, respectively. Working solutions of Y-27632 and CK666 were prepared in media of prescribed viscosity at a 1:1,000 ratio to final concentrations of 10 µM and 100 µM, respectively.

### Software

Single-cell segmentation for measurements of motility and morphological plasticity was conducted using CellPose and CellProfiler^47,48^. Cellular trajectories were tracked using the TrackPy algorithm in Python. All analyses were conducted in Python using these specifications: python 3.11.3, pandas 1.5.3, numpy 1.24.3, trackpy 0.6.2, cv2 4.8.1, plotly 5.9.0, scipy 1.10.1, scikit-learn 1.2.2, seaborn 0.12.2.

### Statistics

Statistical significance for experiments with two groups was calculated using the Mann-Whitney U test or Cohen’s *d* (when *n* ≥ 10,000). Statistical significance for experiments with two groups was calculated using one-way-ANOVA or Cohen’s *d* (when *n* ≥ 10,000). Linear correlations were estimated using the Pearson correlation coefficient (*r*). A *p*-value of less than 0.05 or a Cohen’s *d* value of greater than 0.2 was considered statistically significant.

## Data availability

All data supporting the findings are present within the manuscript and the supplementary documents.

## ACKNOWLEDGMENTS

This work was funded by the following grants: P30 AG021334 (JW, JMP); Catalyst Award—National Academy of Medicine Global Longevity Challenge (JMP, JW, SS); American Federation for Aging Research and Glenn Foundation Junior Faculty Award (JMP); R35 GM157099 (JMP); R01 CA257647 (KK), R01 GM142175 (KK); R35 GM142838 (KMS). We would like to extend special thanks to Dr. Courtney McQueen, Paul Boyle (Biovision Consulting), and Johns Hopkins University’s Editorial Assistance Service Initiative (EASI) for editorial support of the manuscript.

Schematics: Fig. 1a, Fig. 2a, Fig. 4i, Fig. 5a were made using Biorender.

## AUTHOR CONTRIBUTIONS

YD and JMP conceived the study, experimental design, and analysis. YD, AA, AS, ZG, LT, IS, CE, NMi, AF, and KP conducted experiments. YD, CM, NMa, and JMP analyzed data. YD, SS, KK, JW, and JMP interpreted results. YD and JMP wrote initial version of manuscript. YD, KMS, SS, KK, JW, and JMP wrote and edited manuscript. All authors contributed to editing the last version of the manuscript.

## CONFLICT OF INTEREST

YD, AA, KK, and JMP are co-inventors on a patent application related to this study. All other authors declare no conflict of interest.

